# NF2 Loss-of-Function and Hypoxia Drive Radiation Resistance in Grade 2 Meningiomas

**DOI:** 10.1101/2023.09.08.556892

**Authors:** Bhuvic Patel, Sangami Pugazenthi, Collin W. English, Tatenda Mahlokozera, William A. Leidig, Hsiang-Chih Lu, Alicia Yang, Kaleigh Roberts, Patrick DeSouza, Diane D. Mao, Namita Sinha, Joseph E. Ippolito, Sonika Dahiya, Allegra Petti, Hiroko Yano, Tiemo J. Klisch, Akdes S. Harmanci, Akash J. Patel, Albert H. Kim

## Abstract

**Background:** World Health Organization Grade 2 meningiomas (G2Ms) exhibit an aggressive natural history characterized by recurrence and therapy resistance. G2Ms with histopathological necrosis have been associated with worse local control (LC) following radiation therapy, but drivers and biomarkers of radiation resistance in these G2Ms remain unknown.

**Methods:** We performed genetic sequencing and histopathological analysis of 113 G2Ms and investigated the role of intratumoral hypoxia as well as genes of interest through knockdown and clonogenic survival following ionizing radiation. Lastly, we performed transcriptional profiling of our *in vitro* model and 18 G2M tumors using RNA sequencing.

**Results:** *NF2* loss-of-function (LOF) mutations were associated with necrosis in G2Ms (p=0.0127). Tumors with *NF2* mutation and necrosis had worse post-radiation LC compared to *NF2* wildtype tumors without necrosis (p=0.035). Under hypoxic conditions, *NF2* knockdown increased radiation resistance *in vitro* (p<0.001). Bulk RNA sequencing of our *in vitro* model revealed *NF2*- and hypoxia-specific changes and a 50-gene set signature specific to radiation resistant, *NF2* knockdown and hypoxic cells, which could distinguish *NF2* mutant and necrotic patient G2Ms by unsupervised clustering. Gene set enrichment analysis of patient tumor and *in vitro* data revealed downregulation of apoptosis and upregulation of proliferation in *NF2*-deficient and hypoxic cells, which we validated with functional assays.

**Conclusions:** *NF2* LOF in the setting of hypoxia confers radiation resistance through transcriptional programs that reduce apoptosis and promote proliferation. These pathways may identify tumors resistant to radiation and represent therapeutic targets that in the future could improve LC in patients with radiation resistant G2Ms.

**KEY POINTS:** 1. Spontaneous necrosis with *NF2* mutations is associated with radio-resistance in WHO G2Ms.

2. *NF2* knockdown in the setting of hypoxia confers radio-resistance to meningioma cells *in vitro* and is driven by increased cell proliferation and decreased apoptosis.

**IMPORTANCE OF THE STUDY:** World Health Organization Grade 2 meningiomas (G2M) are often treated with surgical resection followed by radiation, especially in the case of recurrence. However, the mechanisms underlying radiation resistance in G2Ms remain to be identified, and moreover, we lack biomarkers to distinguish G2Ms that will respond to radiotherapy from those that are refractory. In this study we perform histological and molecular analysis of a large cohort of G2Ms to identify predictors of radiation resistance. Using these data and an *in vitro* model of radiation therapy, we demonstrate that radiation resistance in G2Ms is likely driven by the combination of *NF2* gene mutations and the hypoxia that accompanies tumor necrosis. Patients whose tumors bear these two features may therefore benefit from alternative treatments that target specific pathways implicated in radiation resistance.

## INTRODUCTION

Meningiomas are the most common primary brain tumor in adults, comprising 39% of all brain tumors and 54.5% of non-malignant brain tumors.^1^ Although most are considered to be benign, a subset of meningiomas present with an aggressive clinical course resistant to treatment. World Health Organization (WHO) Grade 2 meningiomas (G2M),^2^ which account for 20.4-35.5% of meningiomas, are a particularly heterogeneous group of tumors with a wide spectrum of clinical behaviors, making prognostication and treatment more complex.^3^ The current treatment strategy for meningiomas is surgical resection and/or radiation therapy.

Previously published data demonstrated that meningiomas with spontaneous intratumoral necrosis are associated with radiation resistance and an aggressive clinical course marked by frequent recurrence and poor survivorship.^4,5^ Although tumor necrosis may predict radiation resistance, its utility as a marker to identify resistant G2Ms is limited as it relies on accurate identification of necrotic regions of tumor, which may not be present in small tumor specimens and may be overlooked on microscopic examination, especially in large tumors. Discovery of a molecular marker for radiation resistance in G2Ms would enable more reliable identification of resistant tumors, but no such biomarkers have been discovered to date. In addition, the biological role of necrosis in G2M radiation resistance remains unexplored.

In this study, we perform molecular analysis of radiation resistant G2Ms to identify molecular alterations that cause radiation resistance and develop an *in vitro* model to validate our findings We also identify a transcriptional signature that is associated with G2M radiation resistance.

## MATERIALS AND METHODS

### Institutional Approval

All human patient sample sequencing and clinical data analysis was approved by the Institutional Review Board of both institutions from which patient data was gathered.

### Cell culture

IOMM-Lee meningioma cells were grown in Dulbecco’s modified Eagle’s medium with 4.g/L D-glucose supplemented with 10% heat inactivated fetal bovine serum, 1% L-glutamine, 1% sodium pyruvate, 1% nonessential amino acids and 1% penicillin/streptomycin (Life Technologies). Human embryonic kidney (HEK293T) cells were cultured in Dulbecco’s modified Eagle’s medium with 10% heat inactivated fetal bovine serum and 1% penicillin/streptomycin. All cells were incubated at 37 °C and 5% CO_2_.

### Lentiviral transfection

Lentivirus production and transfection were performed as previously described.^6^ Briefly, HEK293T cells were plated with a goal density of 70–80%. Transfection was performed one day later using the PEI transfection method with the experimental plasmid, the packaging plasmid psPAX2, and the envelope plasmid pCMV-VSVG. Four days later, culture medium was collected and centrifuged. The supernatant was then filtered, Lenti-X Concentrator (Clontech) was added, and the mixture was incubated at 4 °C for 6 hours. The solution was then centrifuged at 1500x*g* for 45 min at 4 °C. The supernatant was aspirated, and pellets containing virus were resuspended in one-tenth of the original medium volume of cold PBS and stored at −80 °C. IOMM-Lee cells were infected with *NF2* or SHC002 control lentivirus in 24-well plates, media was changed 6 hours later, and cells underwent puromycin selection two days after infection before being collected two days later (four days after infection).

### Clonogenic Assay

Clonogenic assays were used to evaluate radiation resistance *in vitro* and were performed as previously described.^7^ Briefly, for normoxic condition experiments, cells were plated in Corning 6-well plates at the following concentrations: 1000 cells for 0 Gy, 1500 cells for 2 Gy, 3000 cells for 4 Gy, and 7000 cells for 6 Gy. For hypoxic condition experiments, cell plating densities were doubled and deferoxamine mesylate (DFX, Millipore Sigma catalog #252750) was added to media at 125 μM final concentration 24 hours after plating. Two days after plating, radiation was dosed at 0 Gy, 2 Gy, 4 Gy, and 6 Gy using a Rad Source RS2000 irradiator. The media was changed one day after administration of radiation to remove DFX. For hypoxia chamber experiments, cells were incubated in a O_2_ Control InVitro Glove Box Hypoxia Chamber (Coy Lab Products) instead of adding DFX and were kept in the chamber until one day post radiation except for the short period of time necessary to administer radiation. Five days after media change or removal from the hypoxia chamber, plates were stained with 0.5% crystal violet in 30% methanol and 10% acetic acid. Wells were scanned using brightfield microscope imaging. Images were then randomized and counted by a blinded reviewer. Radiation dose titration curves were modeled using GraphPad PRISM 9 using a linear quadratic cell death model (Fraction of cells surviving = e^-(A*D^ ^+ B*D^2)^) and comparing curves using the extra sum-of-squares F Test.^8^

### EdU Assay

To evaluate cell proliferation post-radiation the Click-iT EdU assay (Thermo Scientific) was used. IOMM-Lee cells were plated in Corning 96 well plates. One day later the media was changed to media containing DFX to create a hypoxic environment. One day later, cells were irradiated at increasing doses (0 Gy, 2 Gy, 4 Gy, 6 Gy). The media was changed one day later to remove DFX and the EdU assay was performed two days thereafter as per the manufacturer protocol. Fluorescent and bright field images of cells were obtained using a Leica DMi8 microscope (Leica Microsystems, Wetzlar, Germany) and were quantified using the Fiji analyze particles function.

### Caspase Assay

Apoptosis was evaluated using the Caspase 3/7 live cell imaging assay. IOMM Lee cells were plated at 1,500 cells per well in Corning 96-well plates, exposed to DFX for 24 hours and then irradiated. 24 hours after irradiation, the Incucyte Caspase-3/7 Green dye (Sartorius) was added at 1:1000 dilution and live cell imaging was performed on the Incucyte S3 System (Sartorius). Apoptosis was calculated using the S3 Live Cell Analysis Instrument (Sartorius) by calculating Green Object Count per Image/Phase Area Confluence.

### Plasmids and Antibodies

The following shRNA plasmids were used: NF2i.1 (TRCN0000039974, Target Sequence: GCTCTGGATATTCTGCACAAT), NF2i.2 (TRCN0000039975, Target Sequence: GCTTCGTGTTAATAAGCTGAT), SHC002 Control (all from Sigma Aldrich, Burlington, MA). Western blot antibodies included: Merlin Rabbit mAb #12888 (Cell Signaling, Danvers, MA) and HIF-1a Mouse mAb #610959 (BD Biosciences, Franklin Lakes, NJ).

### Whole Exome Sequencing

Automated dual indexed FFPE libraries were constructed with 50-250 ng of genomic DNA utilizing the KAPA HTP Library Kit (KAPA Biosystems) on the SciClone NGS instrument (Perkin Elmer) targeting 250 bp inserts. 10 samples were pooled at an equimolar ratio yielding ∼2.2 µg in the pool prior to the hybrid capture. Library pools were hybridized with a probe capture set that consisted of Nimblegen v3 Exome + Regulome + TERT Promoter probes spanning a 125Mb target region of the human genome. KAPA qPCR was used to quantify the libraries and determine the appropriate concentration to produce optimal recommended cluster density on a HiSeq4000 v2 (PE150bp) sequencing run. All sequencing was completed according to the manufacturer’s recommendations (Illumina Inc, San Diego, CA). Alignment of sequencing reads to the human reference genome grch37, variant calling and copy number analysis were performed using bedtools (v2.17.0), picard (v1.113), bwamem (v0.7.10), samtools (v0.1.19), varscan (v2.3.6), unionunique, and CNVkit.

### RNA Sequencing

Dual indexed FFPE libraries were constructed with a 500 ng input of total RNA utilizing the TruSeq Stranded Total RNA Library Construction riboGold kit (Illumina). Two pools of five samples each were pooled at an equimolar ratio yielding ∼900ng (Pool 1) −1600 ng (Pool 2) in the pool prior to the hybrid capture. Library pools were hybridized with the xGen Exome Research Panel v1.0 reagent (IDT Technologies) that spans 39 Mb target region (19,396 genes) of the human genome. KAPA qPCR was used to quantify the libraries and determine the appropriate concentration to produce optimal recommended cluster density on a HiSeq4000 v2 (PE150bp) sequencing run. All sequencing runs were completed according to the manufacturer’s recommendations (Illumina Inc, San Diego, CA).

### RNA Sequencing Analysis

All gene-level and transcript counts were imported into the R/Bioconductor package EdgeR and TMM normalization size factors were calculated to adjust for differences in library size. Genes or transcripts not expressed in any sample were excluded from further analysis. The TMM size factors and matrix of counts were imported into R/Bioconductor package limma, and weighted likelihoods based on the observed mean–variance relationship of every gene/transcript were calculated for all samples with the Voom function. The performance of the samples was assessed with a Spearman correlation matrix and multidimensional scaling plots. Gene/transcript performance were assessed with plots of residual SD of every gene to their average log-count with a robustly fitted trend line of the residuals. Generalized linear models with robust dispersion estimates were created to test for gene/transcript-level differential expression. Differentially expressed genes (DEGs) and transcripts between sample groups were identified. Unsupervised clustering was performed using Spearman’s method and heatmap generation was performed using ComplexHeatmap R package (Version 2.16.0). Gene set enrichment analysis (GSEA) was performed using the gene ontology (GO) biological process (BP) gene set in the experimental group compared to control using fsgsea R package (1.26.0).

### Statistical analysis

Chi square analysis with Fisher’s exact test was performed to evaluate associations between necrosis status and *NF2* mutational status, 22q chromosomal loss status, and 1p chromosomal loss status. Log rank test was used to analyze local control curves. GraphPad PRISM 9 was used to perform statistical analyses.

## RESULTS

### NF2 mutations and intratumoral necrosis are associated with radiation resistance

To discover any previously undescribed mutations associated with radiation resistance, we first performed whole exome sequencing on a discovery cohort of ten G2Ms, five with intratumoral necrosis and recurrence after radiation therapy (*i.e.,* radiation resistant tumors) and five without intratumoral necrosis and which responded to radiation therapy without recurrence (*i.e.,* radiosensitive tumors). Since necrosis has previously been associated with radiation resistance,^4^ DNA was extracted from an approximately 3 mm wide “ring” of tissue surrounding the necrotic region to maximize detection of possible drivers of radiation resistance (Figure 1A). Demographic information and survival data for tumors included in each of the two groups are shown in Figure 1B and 1C. As expected, meningioma-associated mutations previously described in the literature were present in the discovery cohort with the exception of the *TERT* promoter mutation and CDKN2A/B loss, which are now considered a molecular marker of WHO grade 3 meningiomas (Figure 1D).^2^ Since matched normal DNA from subjects was not available to filter germline variants, only non-synonymous variants in genes appearing in the COSMIC database or previously described in meningiomas in the literature were included for further analysis.^9,10^ In total, nine recurrently mutated genes were identified in each group. Enrichment of recurrent mutations in each group was statistically assessed using a Fisher’s exact test, and only mutations in the *NF2* gene were found to be significantly enriched in the radiation resistant group after correction for multiple comparisons (p = 0.028, Benjamini-Hochberg false discovery rate 0.25).

**Figure 1.**
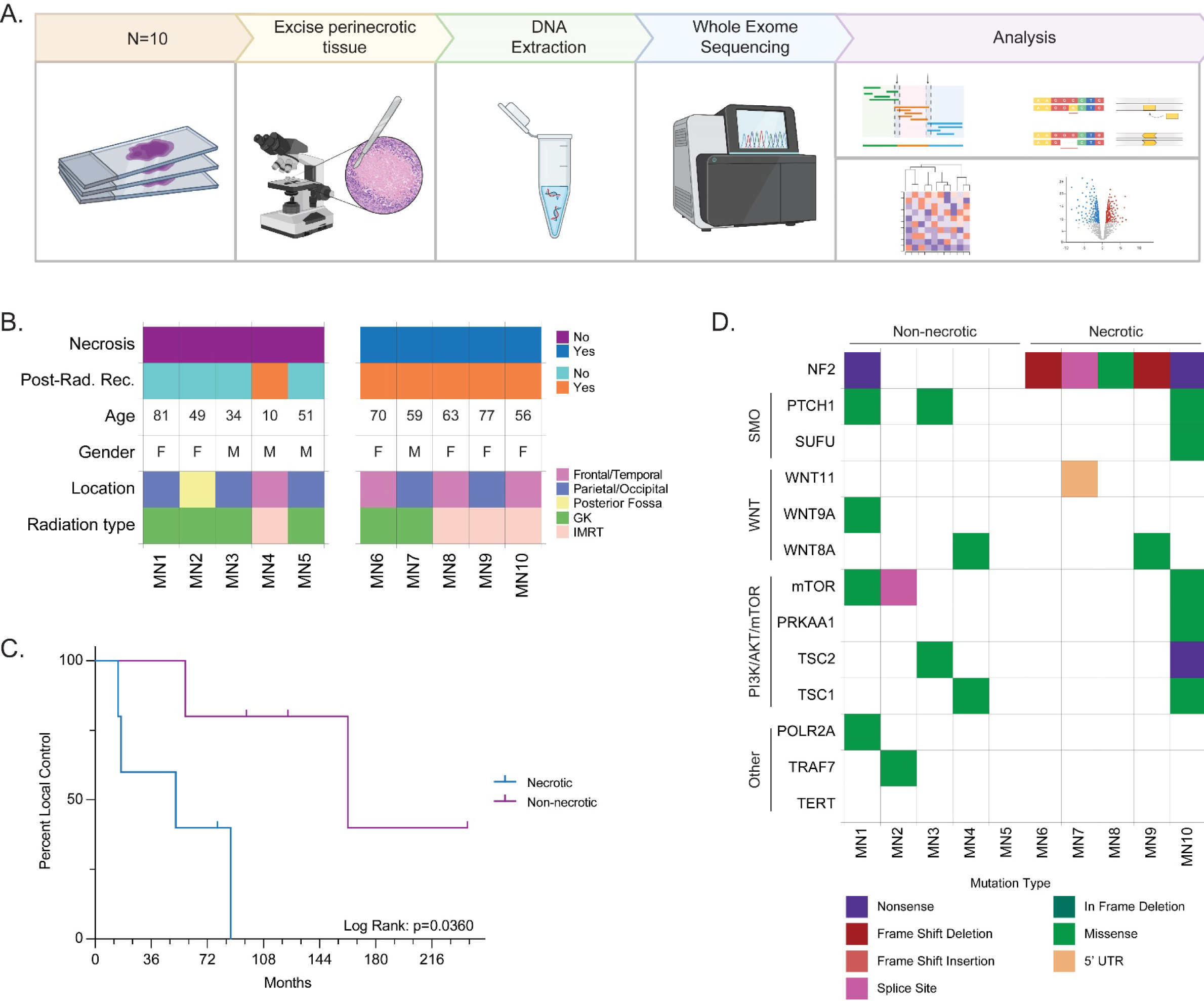
Whole exome sequencing of discovery cohort of G2M. (A) Workflow schematic for whole exome sequencing of G2Ms in discovery cohort (n=10). (B) Discovery cohort patient and clinical characteristics. (C) Local control curve between G2M with and without spontaneous necrosis (n=5, each). (D) Presence of nonsynonymous variants of genes from COSMIC database or previously described in meningiomas in necrotic vs nonnecrotic G2Ms. *p=0.024

To validate the results from the discovery cohort, targeted panel sequencing was performed on a larger cohort of G2M. The validation cohort was comprised of 93 G2Ms from two institutions, 76 from one institution and 17 from another (Figure 2A). Histological evaluation and targeted sequencing of these tumors also demonstrated associations between spontaneous necrosis and *NF2* mutations (Fisher’s Exact: p=0.0127; Figure 2B), corroborating our discovery cohort results. 1p and 22q loss were not significantly associated with necrosis in our validation cohort, highlighting the particular importance of *NF2* loss of function mutation (Figure 2B).

**Figure 2.**
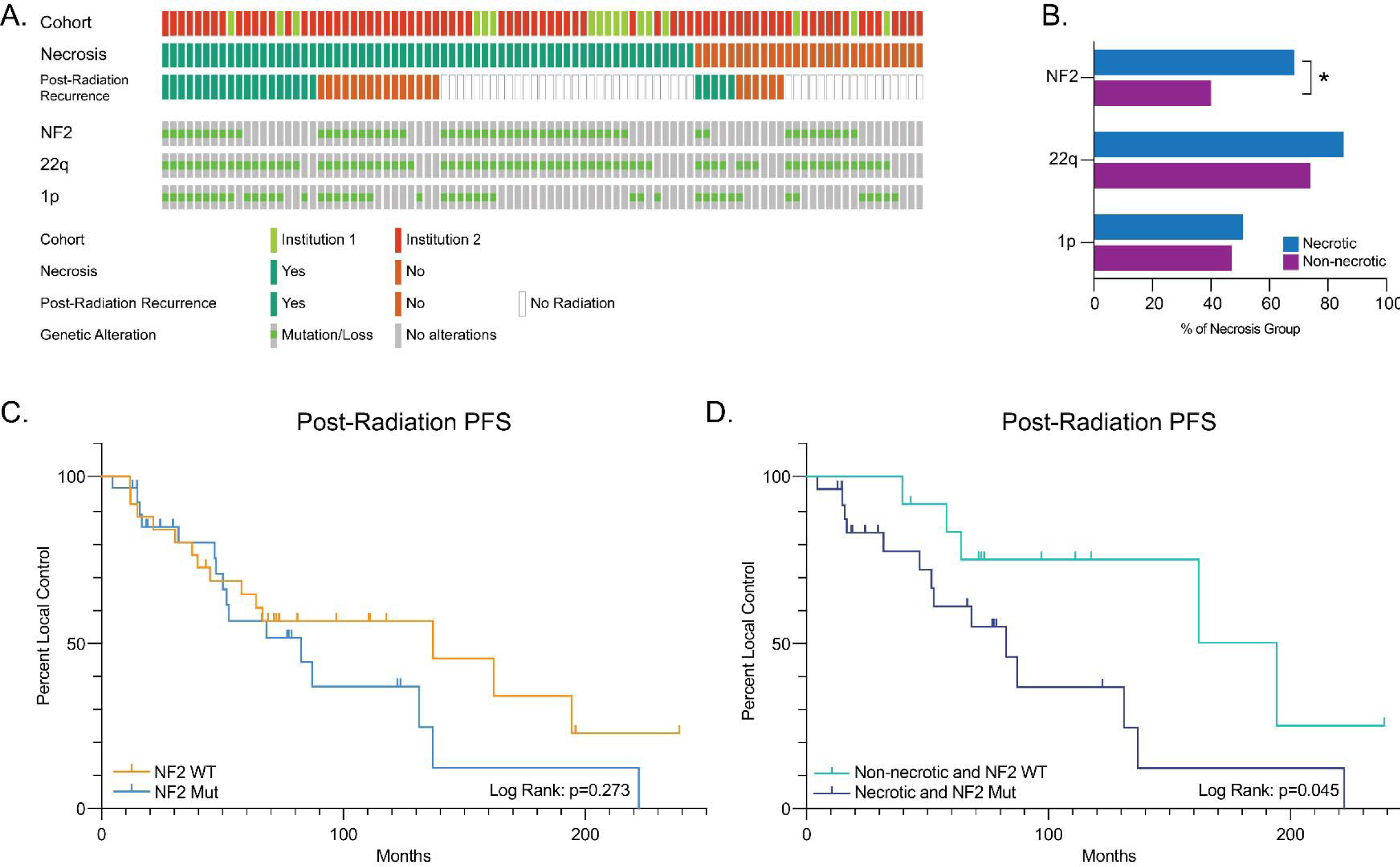
Mutational and histopathological profiling of large validation cohort of G2M. (A) Oncoprint of validation cohort of G2M from two institutions (n=103. Necrotic status, post radiation recurrence, NF2 mutation, 22q loss, and 1p loss is represented. (B) Percent of necrotic or non-necrotic group with NF2 mutation, 22q loss, or 1p loss. *p=0.0127. (C) Percent local control curve comparing post-radiation recurrence of G2Ms without (n=26) and with (n=29) NF2 mutation. (D) Percent local control curve comparing post-radiation recurrence of non-necrotic G2Ms without NF2 mutation (n=13) against necrotic G2Ms with NF2 mutation (n=26).

Next, we tested the hypothesis that G2Ms with spontaneous necrosis and *NF2* mutation have increased radiation resistance by evaluating local control after radiotherapy for all tumors treated with radiotherapy (n = 45) using survival data from these patients. We first compared tumors with and without *NF2* mutations, without considering intratumoral necrosis and found that *NF2* mutational status alone was not associated with decreased local control (Log Rank: p=0.273; Figure 2C). We then compared tumors with and without necrosis, without considering *NF2* mutational status, and found that histopathological necrosis alone was also not associated with decreased local control, although a trend towards association was observed (Log Rank: p=0.057; Supplemental Figure 1). Lastly, we considered the possibility that the combination of *NF2* mutational status and histopathological necrosis may be associated with decreased local control. Interestingly, we found that patients with both spontaneous necrosis and *NF2* mutation demonstrated significantly worse local control than those without spontaneous necrosis and *NF2* mutation (Log Rank: p=0.045; Figure 2D), suggesting that the combination of necrosis and *NF2* mutation contribute to radiation resistance in G2Ms.

### Modeling radiation in vitro uncovers a critical role for NF2 loss of function and concurrent hypoxia in radiation resistance

To investigate the mechanisms of radiation resistance in G2Ms, we developed an *in vitro* model of radiation response using the IOMM-Lee meningioma cell line, which has wildtype *NF2* alleles, thus enabling manipulation of NF2 expression (Figure 3A). To model *NF2* loss of function mutations, we used lentiviral RNA interference to target *NF2* and verified robust knockdown of the NF2 protein Merlin by immunoblot (Figure 3B). Prior work using Crispr/Cas *NF2* knockdown in IOMM-Lee meningioma cells suggests that they form larger colonies compared to WT cells, leading to our hypothesis that *NF2* knockdown may also confer radiation resistance.^11^ To evaluate radiation resistance in meningioma cells, we performed clonogenic assays following exposure to increasing doses of radiation. Under normoxic conditions, we observed a small but significant difference in clonogenic survival with only one of the two *NF2* shRNAs (*NF2i.1*) following ionizing radiation (IR) compared to control shRNA (*NF2i.1*: p<0.0001, F = 10.84; *NF2i.2:* p=0.1632, F = 1.988) (Figure 3C), overall suggesting a modest, if any, effect of *NF2* loss by itself on radiation resistance.

**Figure 3.**
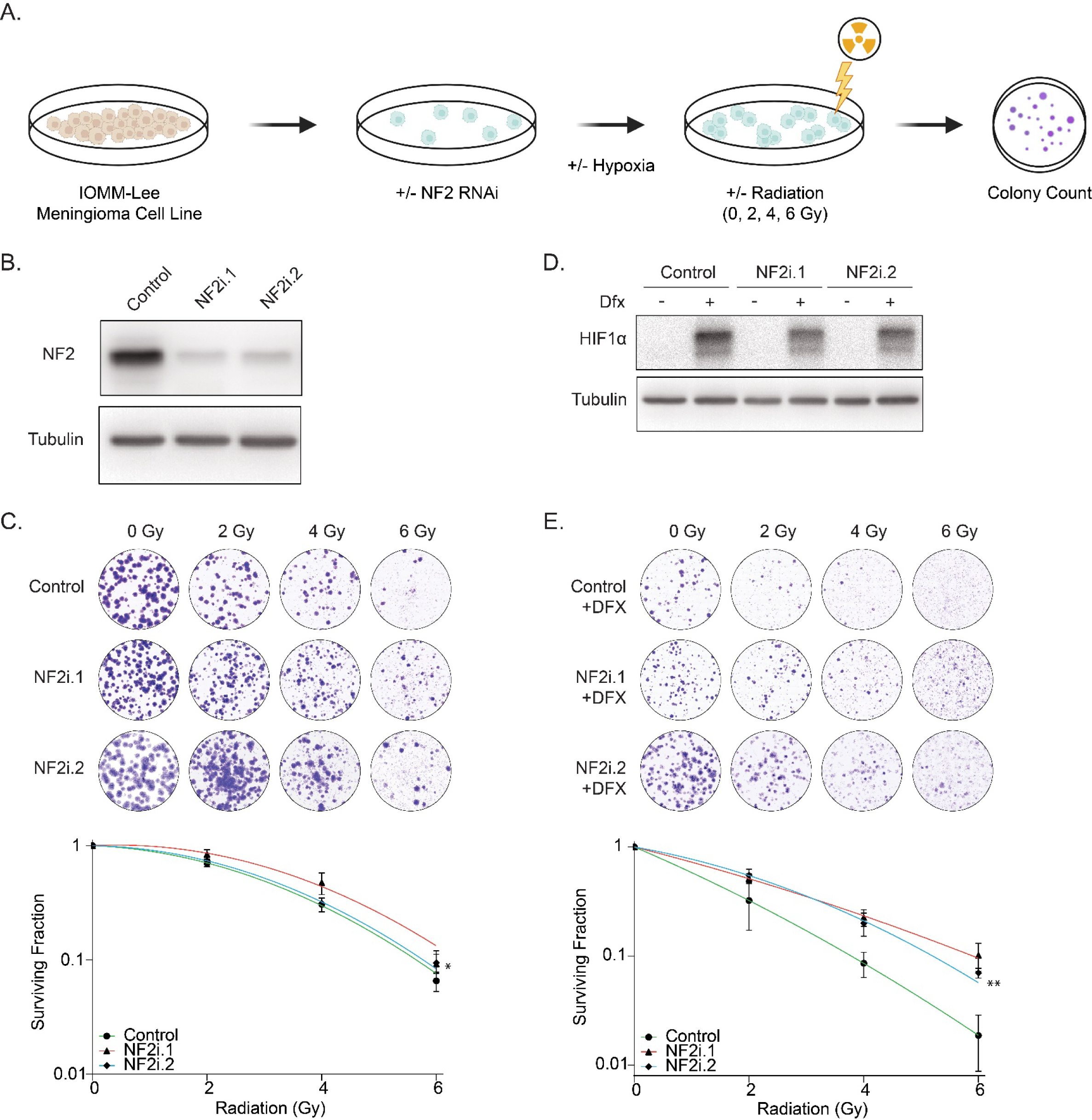
*In vitro* model of radiation resistance. (A) Schematic depicting *in vitro* model using lentivirus infection, radiation exposure, and clonogenic assay. (B) Western blot showing NF2 knockdown of NF2i.1 and NF2i.2 lines based on Merlin protein concentration. (C) Western blot showing induction of HIF1a from DFX induced hypoxia. (D) Clonogenic assay with NF2i lines in normoxia with representative colony images with increasing doses of radiation. Surviving fraction of colonies for control, NF2i.1 and NF2i.2 lines with increasing doses of radiation. *NF2i.1* vs. control: p<0.0001, F = 10.84; *NF2i.2* vs. control: p=0.1632, F = 1.988. (E) Clonogenic assay with NF2i lines in hypoxia with representative colony images with increasing doses of radiation. Surviving fraction of colonies for control, NF2i.1 and NF2i.2 lines in hypoxia with increasing doses of radiation. *NF2i.1* vs. control: p<0.0001, F=16.22; *NF2i.2* vs. control: p<0.0001, F=16.07.

Based on our clinical data, which showed worse progression free survival in tumors with combined *NF2* mutation and necrosis, we hypothesized that combined *NF2* knockdown and hypoxia might confer more robust radiation resistance to meningioma cells. Intratumoral necrosis has previously been shown to be driven at least in part by intratumoral hypoxia.^12,13^ We therefore chemically induced hypoxia in our cells using deferoxamine mesylate (DFX), a chelator of iron oxide that reduces steady-state dissolved oxygen in culture media.^14,15^ As expected, cells grown in media containing DFX had increased expression of HIF1α, a well-established marker of hypoxia response, by immunblot (Figure 3D).^16^ Under hypoxic conditions, we observed a robust increase in clonogenic survival following IR in *NF2*-knockdown cells, using two distinct RNAi, compared to control cells (*NF2i.1*: p<0.0001, F=16.22; *NF2i.2*: p<0.0001, F=16.07; Figure 3E). These clonogenic survival results were recapitulated using a hypoxia chamber instead of DFX. (Supplemental Figure 2). Thus, we found that rather than *NF2* loss alone, hypoxia and *NF2* knockdown together confer radiation resistance to meningioma cells *in vitro*.

### Analysis of in vitro gene expression in radiation resistant meningioma cells reveals transcriptional contributions of NF2 loss and hypoxia

To elucidate the roles of *NF2* knockdown and hypoxia in radiation resistance, we used bulk RNA sequencing to perform transcriptional profiling of cells in our model system. First, to identify hypoxia-driven transcriptional alterations, we compared gene expression profiles of WT cells in hypoxia to WT cells in normoxia. As expected, the top DEGs included those previously shown to be related to hypoxia response mechanisms such as *PTGER2*, *CMPK2*, and *PLK2*.^17–19^ Next, to identify transcriptional changes related to *NF2* loss of function, we compared *NF2* knockdown cells in hypoxia to WT cells in hypoxia. This comparison yielded a list of DEGs including the following that have been previously implicated in meningioma biology or other cancers: *CALCRL, CDH1*, *SDC2*, *PLAC8*, and *CFH*.^20–25^ Following analysis of DEGs driven by hypoxia or *NF2* knockdown, we sought to identify transcriptional changes attributable to *NF2* knockdown in the context of hypoxia, which induced radiation resistance in our *in vitro* model. To do so, we compared the transcriptional profile of the radiation resistant line (*NF2* knockdown cells in hypoxia) to the radio-sensitive line (WT cells in normoxia). When comparing the top 25 upregulated and downregulated DEGs associated with radiation resistance to the DEGs associated with *NF2* knockdown or hypoxia exposure alone, we found that the upregulation of *IP6K3*, *ASPG*, *SPANXA1*, *SPANXA2* and downregulation of *JAKMIP2* and *FAM107* were shared with *NF2* knockdown alone, but no DEGs were shared with hypoxia exposure alone, suggesting that the predominant contribution to radiation resistance is from *NF2* loss of function. Interestingly, these *NF2*-loss dependent transcriptional alterations have not been previously implicated in *NF2* pathophysiology and may represent novel targets for further investigation and treatment of radiation resistant tumors. Indeed, some of these genes have been implicated in radiation response in other cancers. For example, inhibition of *FAM1315B*, which is upregulated in IOMM-Lee radiation resistant cells *in vitro*, has been previously shown to enhance radiosensitivity in esophageal squamous cell carcinoma, making it a potential target for sensitization of G2Ms to radiation.^26^

### Integration of in vitro and patient G2M RNA sequencing data yields a radiation resistance specific transcriptional signature

Next, we investigated whether expression patterns observed in our *in vitro* model system could be used to differentiate patient tumors with *NF2* mutations and necrosis, a marker of radiation resistance, from *NF2* WT and non-necrotic tumors. We began with a gene set of 200 DEGs (top 100 upregulated and top 100 downregulated by fold change) from the *in vitro* dataset — *NF2* knockdown cells in hypoxia *vs* control cells in normoxia. However, this gene set was unable to reliably separate the *in vivo* tumors into *NF2* and necrotic *vs* WT and non-necrotic groups. The DEG list was thus systematically filtered until we arrived at a gene set of 50 DEGs (top 25 upregulated and top 25 downregulated) from the *in vitro* comparison, which we found could accurately stratify the *in vivo* tumors into two groups (*NF2* and necrotic *vs* WT and non-necrotic) using unsupervised clustering (Figure 4A). These findings demonstrate the generalizability of our *in vitro* model to human G2Ms and highlight the potential utility of these genes as a signature for predicting post-radiation recurrence in G2Ms.

**Figure 4.**
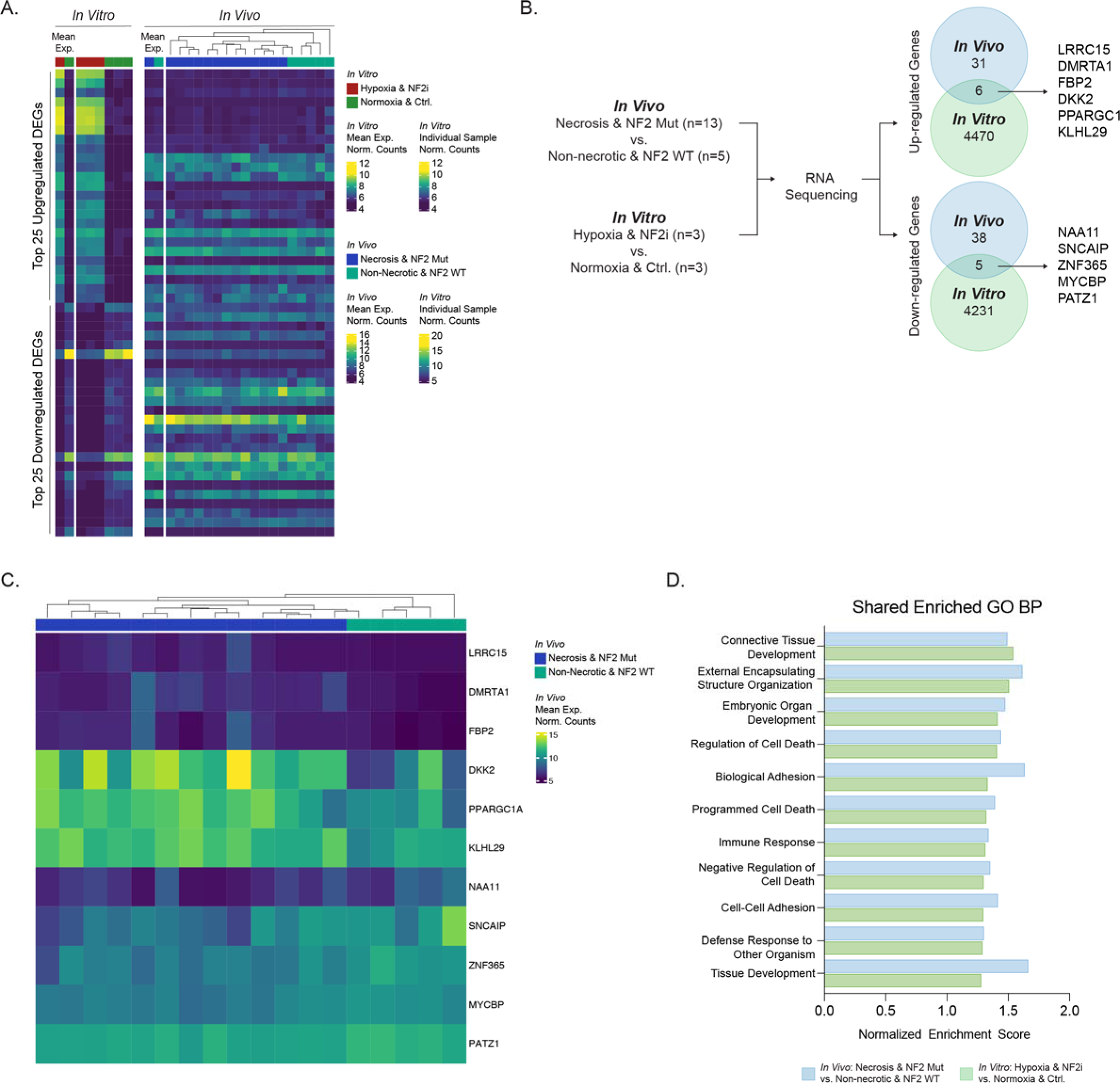
Bulk RNA sequencing analysis of *in vivo* and *in vitro* datasets. (A) Heatmap showing top 25 up and top 25 down regulated DEGs in the *in vitro* samples (*NF2* knockdown in hypoxia *vs NF2* WT in normoxia). Unsupervised clustering of *in vivo* samples (*NF2* mutation and necrotic *vs NF2* WT and non-necrotic) using the 50 DEGs identified from the *in vitro* data. (B) Diagram showing bulk RNA sequencing of *in vivo* and *in vitro* datasets, including the number of common up and down regulated DEGs. *In vivo,* necrotic and NF2 mutant tumors were compared to non-necrotic and NF2 WT tumors. *In vitro,* NF2i cells in hypoxia were compared to control cells in normoxia. (C) Unsupervised clustered heatmap of in vivo samples used 11 shared DEGs identified in (B). (D) Shared enriched GO Biological processes between *in vivo* and *in vitro* datasets described in (B).

To understand the transcriptional effect of radiation on the radiation resistant cells (*NF2* knockdown in hypoxia), we performed DEG analysis comparing *NF2* knockdown cells in hypoxia with and without exposure to 4 Gy radiation. This identified 28 DEGs associated with radiating the radiation resistant cells, of which 21 were upregulated and 7 were downregulated after radiation. Of the 21 upregulated DEGs, *RTL9* and *H2BC4* were upregulated even before radiation exposure in the radiation resistant cells compared to control. Of the 7 downregulated DEGs, *RRM2*, *CCNE2*, and *FAM111B* were significantly downregulated prior to radiation in comparison to control cells. Interestingly, *CCNE2* has been previously shown to be downregulated in the presence of radiation.^27^

To further narrow down the transcriptional changes most likely to be driving radiation resistance in G2Ms we integrated both the human tumor and *in vitro* data by comparing necrotic, *NF2* mutant patient tumors to non-necrotic, *NF2* WT tumors (*in vivo* comparison) and *NF2* knockdown IOMM Lee cells in hypoxic conditions to control cells in normoxic conditions (*in vitro* comparison). When analyzing the upregulated and downregulated DEGs by mean expression for both the *in vivo* and *in vitro* comparisons, we found 6 shared upregulated DEGs (*LRRC15, DMRTA1, FBP2, DKK2, PPARGC1A, KLHL29*) and 5 shared downregulated DEGs (*NAA11, SNCAIP, ZNF365, MYCBP, PATZ1*) between the two datasets (Figure 4B). Interestingly, *DKK2* has previously been shown to be upregulated in the context of radiation resistance in esophageal cancer^28^ and has been associated with more aggressive behavior, such as tumor immune evasion in colorectal cancer^29^ and Ewing sarcoma invasiveness.^30^ When this 11-gene set (6 shared upregulated and 5 shared downregulated DEGs) was used to perform unsupervised clustering of the patient G2Ms, it separated tumors into *NF2* mutation and necrotic *vs NF2* WT and non-necrotic groups (Figure 4C). These results further demonstrate how our *in vitro* model recapitulates the molecular biology of patient tumors and identify several genes that warrant further investigation as both biomarkers and therapeutic targets.

### RNA-sequencing and functional assays implicate increased cell proliferation and decreased cell death in radiation resistance

To identify biological processes potentially responsible for driving radiation resistance, we performed gene set enrichment analysis (GSEA) to identify enriched expression of the gene ontology biological processes (GO BP) gene sets in our RNA sequencing data from the *in vivo* and *in vitro* comparison sets. Relative gene expression values were derived from the comparisons highlighted in Figure 4B (*NF2* knockdown in hypoxia *vs NF2* WT in normoxia (*in vitro* comparison), *NF2* mutation and necrosis *vs NF2* WT and non-necrotic (*in vivo* comparison) and used for GSEA. Shared enriched GO BP terms were then identified between both *in vitro* and *in vivo* comparisons. In total, 11 shared terms were identified, including several notable terms such as programmed cell death, positive regulation of cell death, and negative regulation of cell death (Figure 4D). Cell death associated genes such as caspases (*CASP1* and *CASP8*) as well as *HOXA5*, and *USP17L15* were upregulated in the *in vitro* comparison of *NF2* knockdown cells in hypoxia compared to WT cells. Additionally, inspection of enriched terms not shared between the two comparisons was notable for several terms related to cell proliferation characterized by upregulation of *EGR1*, *SERPINB7*, *CFLAR*, and *PDGFB* (Supplemental Figure 3). Taken together, these data suggested a possible role of regulation of cell death, cell proliferation, or both in radiation resistance.

To further investigate the role of these two processes in radiation resistance, we performed functional assays to measure cell proliferation and cell death using our *in vitro* model system. EdU assays were used to evaluate cell proliferation 72 hours post-radiation in NF2 knockdown and hypoxic (*i.e.* radiation resistant) IOMM-Lee cells. In the absence of radiation, there was no difference in proliferation between the radiation resistant and control meningioma cells. However, following 4 Gy of radiation, *NF2* knockdown cells in hypoxic conditions had a higher proportion of proliferating cells compared to control cells (Figure 5A-B). We next used a caspase-3/7 assay to measure radiation-induced apoptosis in IOMM-Lee cells under hypoxic conditions (Figure 5C). When comparing area under the curve measurements for caspase 3/7 activity, there was no significant difference in apoptosis between WT and *NF2* knockdown IOMM Lee cells without radiation (p=0.2958; Figure 5D). However, after 4 Gy of radiation, the *NF2* knockdown IOMM-Lee cells had significantly lower levels of apoptosis compared to control IOMM Lee cells (p=0.0030; Figure 5D). Thus, radiation resistant G2M cells exhibit increased cell proliferation and decreased apoptosis compared to control cells when exposed to ionizing radiation.

**Figure 5.**
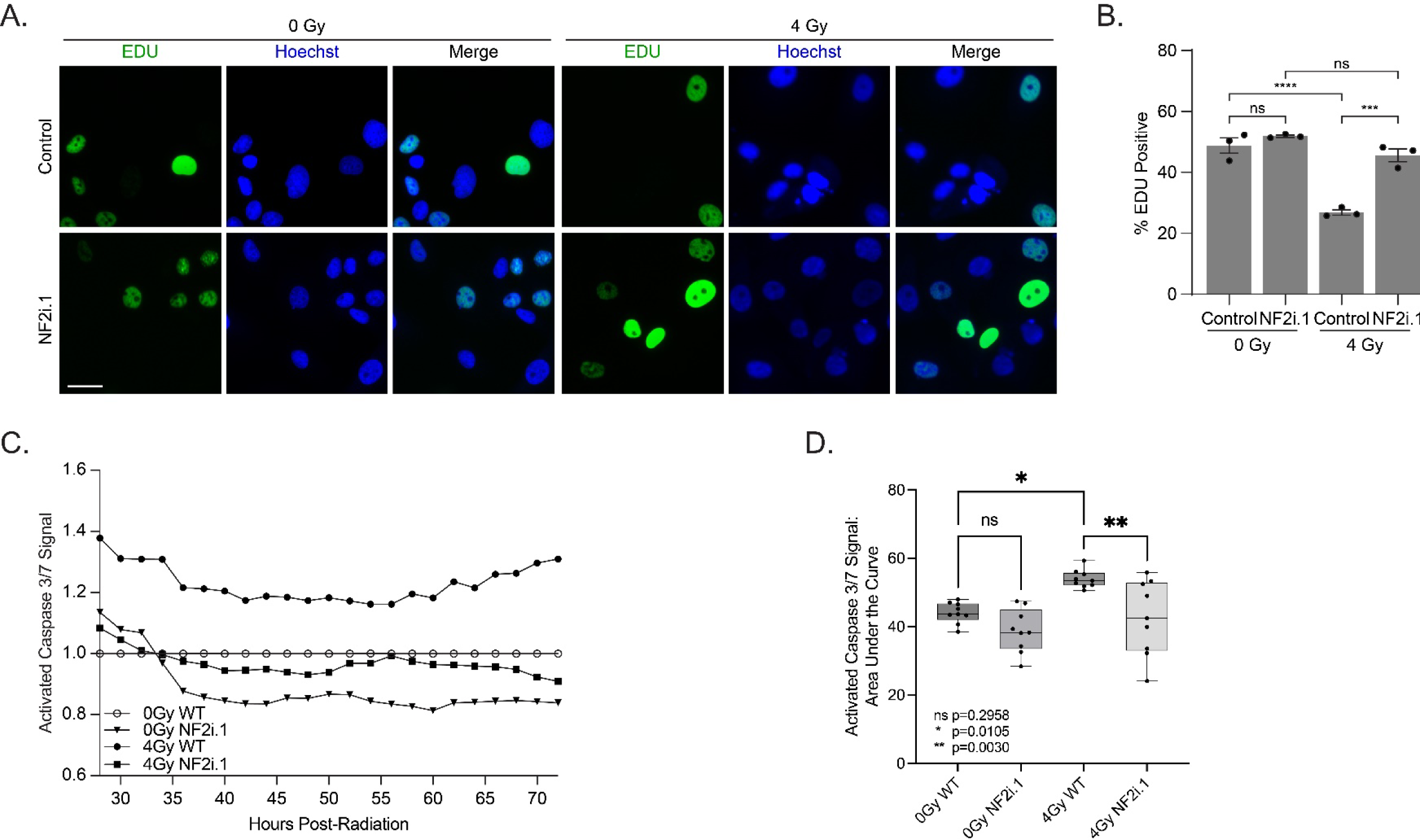
*In vitro* validation of biological processes driving radiation resistance. (A) EdU assay microscopy images for IOMM Lee control and NF2i.1 lines. Scale bar is 20 μm and applies to all panels. (B) Quantification of percent of positive conditions for each condition normalized to cell number based on Hoechst stain. ns: p>0.05, ***p=0.0003, ****p=0.0001. (C) Number of apoptotic cells (defined by Caspase 3/7 signal) normalized to phase area confluence of IOMM Lee control and NF2i.1 cells at 0 Gy and 4 Gy from 28-72 hours post radiation. (D) Area under the curve calculation from (C). ns: 0.2958, *p=0.0105, **=p=0.00

## DISCUSSION

This study highlights discoveries important for both identification and treatment of radiation resistant WHO G2M. Clinical data and targeted/whole exome sequencing analysis of a discovery and validation cohorts demonstrated that spontaneous necrosis on histopathology and *NF2* mutation were associated with post-radiation recurrence in G2Ms. *In vitro* experiments revealed that under hypoxic conditions, IOMM Lee meningioma cells with *NF2* knockdown had increased resistance to radiation compared to WT cells. Bulk RNA sequencing was used to create a 50-gene signature that was able to separate *NF2* mutant and necrotic (radiation resistant) from NF2 wildtype and non-necrotic (radio-sensitive) patient tumors. Lastly, RNA sequencing and functional assays suggested that radiation resistance may be conferred at least in part by increased cell proliferation and decreased apoptosis.

Poor prognostic factors for meningioma recurrence following treatment are older age, male gender, and no or subtotal surgical resection.^31^ Imaging findings associated with meningioma recurrence include lack of focal or diffuse calcification, peritumoral edema, greater tumor volume, and close proximity to a major sinus.^32^ From a histopathological perspective, brain invasion and high mitotic index have been associated with increased recurrence risk and shorter disease-free survival, respectively.^33,34^ We previously reported an association between spontaneous necrosis and G2M radiation resistance following subtotal resection and radiation exposure,^4^ and others have shown that higher MIB1-index is associated with worse local control in G2Ms following radiation therapy.^35^ In this study, we now show that histopathological necrosis and *NF2* mutation together are a strong predictor for radiation resistance in G2Ms.

Although other studies have reported that the presence of spontaneous necrosis in G2Ms predicts worse PFS,^36^ none have validated our previous findings regarding radiation resistance or explored the molecular pathophysiology of spontaneous necrosis in G2Ms. The role of *NF2* mutation, however, has been generally investigated in meningioma prognosis particularly in the context of molecular classification of meningiomas.^37^ Early work suggested that meningiomas harboring *NF2* mutations were more aggressive lesions occurring in convexity, parasagittal, and falcine locations, in contrast to non-*NF2* mutant meningiomas occurring in the skull base.^10,38^ *NF2* mutations have been reported to be the most common mutations in primary G2Ms, with co-occurrence of other aggressive molecular features such as genomic instability or SMARCB1 mutations.^39^ Although we did not find that *NF2* mutation alone was a worse predictor of local control in G2Ms, recent work in a small cohort of 19 grade 2 and 3 meningiomas found that *NF2* WT patients had a 78% reduction in the risk of recurrence following radiation treatment compared to *NF2* mutant patients.^40^ Independently, both spontaneous necrosis and *NF2* mutation have been implicated in aggressive meningiomas, but they have not been discussed together in the context of radiation resistance. Our data suggests that the co-occurrence of these features in meningiomas is a troubling sign regarding radio-sensitivity and local control.

Molecular analysis of meningiomas has been predominantly focused on identifying markers for recurrence. Many meningiomas exhibit both a loss of heterozygosity on chromosome 22 as well an inactivating *NF2* mutation.^41^ Higher grade tumors are more likely to have additional chromosomal abnormalities, loss of function (LOF) *NF2* mutations, and a higher overall mutational burden.^42^ Several other molecular characteristics have been implicated in meningioma recurrence such as increased Ki-67 proliferation index and increased expression of human epidermal growth factor receptor 2 (HER2) protein, proliferating cell nuclear antigen (PCNA), and telomerase reverse transcriptase (*TERT*).^43^ Driver mutations in *NF2*, phosphoinositide 3-kinase (PI3K, specifically in *PIK3CA*), sonic hedgehog pathway, *TERT* promoter, and *tumor necrosis factor receptor-associated factor 7* (*TRAF7*) have also been identified as more common in recurrent meningiomas.^44^ Due to the inadequacy of WHO grade, clinical characteristics, or DNA mutations alone in predicting meningioma prognosis, some groups have used a combination of molecular techniques and clinical course to create classification schema that more accurately predict meningioma outcomes.^45–48^ The importance of molecular markers predicting meningioma clinical course was most recently reflected in the 2021 WHO grading criteria, which include both the *TERT* promotor mutation and *CDKN2A/B* homozygous loss as markers of Grade 3 meningiomas.^2^ Notably, it has been shown that *CDKN2A* mutations almost exclusively occur in *NF2* mutated meningiomas.^49^ Following our work and in conjunction with prior investigations, there may be merit to including *NF2* mutation into the WHO grading criteria especially in combination with histopathological spontaneous necrosis as a marker of a more aggressive lesion resistant to radiation.

Additionally, several recent studies have investigated molecular classifications of meningiomas that predict meningioma clinical behavior more accurately than the existing WHO criteria.^45,47,48^ Together, these studies identified three major meningioma molecular classes (Group A/Merlin intact, Group B/Immune enriched/, and Group C/Hypermitotic), with the most benign class having wildtype *NF2* alleles. The two more aggressive classes both are defined partially by *NF2* loss, indicating that likely radiation resistant meningiomas are in the two more aggressive molecular classes of meningiomas.^50^ Further mechanistic evaluation of the three major molecular classes showed that the Merlin rescue in CH-157MN xenografts increased apoptosis in response to ionizing radiation compared to meningiomas without Merlin.^50^ This murine data corroborates our *in vitro* data, which shows that in the context of *NF2* knockdown, meningiomas have lower levels of apoptosis when exposed to radiation.

Through molecular analysis, we identified several gene targets that may have utility in sensitizing radiation resistant G2Ms to radiation. For example, *DKK2*, which is involved in the Wnt/β-catenin signaling pathway, was upregulated in both *in vitro* radiation resistant cells and radiation resistant *in vivo* G2Ms and has been shown to be upregulated in radiation resistant esophageal cancer^28^ and has been associated with more aggressive behavior in other cancers. ^29,30^ Additionally, *FAM1315B*, which is upregulated in radiation resistant cells *in vitro*, has been previously shown to enhance radiosensitivity when inhibited in esophageal squamous cell carcinoma.^26^ *CCNE2,* a gene downregulated when radiation resistant cells were exposed to radiation, has been previously shown to be downregulated in the presence of radiation as well.^27^ Thus, the results of our analysis and supporting evidence from previous studies may indicate that these genes may represent potential targets to sensitize G2Ms to radiation.

Although this work is strengthened by being multi-institutional and including both *in vitro* and *in vivo* data, it is not without limitations. Importantly, because it is focused on WHO G2Ms, these results may not be generalizable to WHO grade 1 or grade 3 tumors. Additionally, our analysis is limited to patients who underwent surgical resection of their tumors and relies primarily on targeted panel sequencing which leaves open the possibility of yet-undiscovered genomic alterations driving radiation resistance. Lastly, our proposed gene signature for radiation resistance will require independent validation in a separate cohort of G2Ms in future studies.

## CONCLUSIONS

We found that spontaneous necrosis and *NF2* mutation in patients with G2Ms were associated with radiation resistance. *In vitro* validation revealed that meningioma cells with *NF2* knockdown in the presence of hypoxia were radiation resistant compared to control cells. This phenotype is in part due to increased apoptosis and decreased cell proliferation of the radiation resistant group. Overall, this study has significant implications both for prognostic evaluation of G2M patients and our understanding of the biology of radiation resistance in G2Ms.

## FUNDING

**BP:** 2019-2020 Neurosurgery Research and Education Fund and Academy of Neurological Surgeons Research Fellowship Grant

**SP:** Washington University School of Medicine Dean’s Medical Student Research Fellowship for the MD5 Yearlong Research Program; 2023 NREF Medical Student Summer Research Fellowship; American Brain Tumor Association Jack & Fay Netchin Medical Student Summer Fellowship in Honor of Paul Fabbri

**AHK:** Alvin J. Siteman Cancer Research Fund (GF0010215), the Christopher Davidson and Knight Family Fund, and the Duesenberg Research Fund

## CONFLICT OF INTEREST

A.H.K. is a consultant for Monteris Medical and has received non-related research grants from Stryker for study of a dural substitute.

## AUTHORSHIP

Project conception: BP, AHK

Data collection: BP, SP, AY, CWE

Data analysis: BP, SP

Manuscript drafting: BP, SP, AHK

Manuscript review: AJP, AHK

Manuscript approval: All

## DATA AVAILABILITY

Data will be made available upon reasonable request.

## ACKNOWLEDGMENTS

We would like to acknowledge: Gavin Dunn and his laboratory for providing IOMM Lee cell lines and cell culture methods, and Hiroko Yano Lab and Albert Kim Lab members for their support on this project with *in vitro* experiments.

## PREVIOUS PRESENTATION

Portions of this work were presented at the 2021 Society for Neuro-Oncology Annual Meeting and the 2021 and 2023 Congress of Neurological Surgeons Annual Meetings.

