## Supplementary Figures for "NF2 Loss-of-Function and Hypoxia Drive Radiation Resistance in Grade 2 Meningiomas"

Supplemental Figure 1

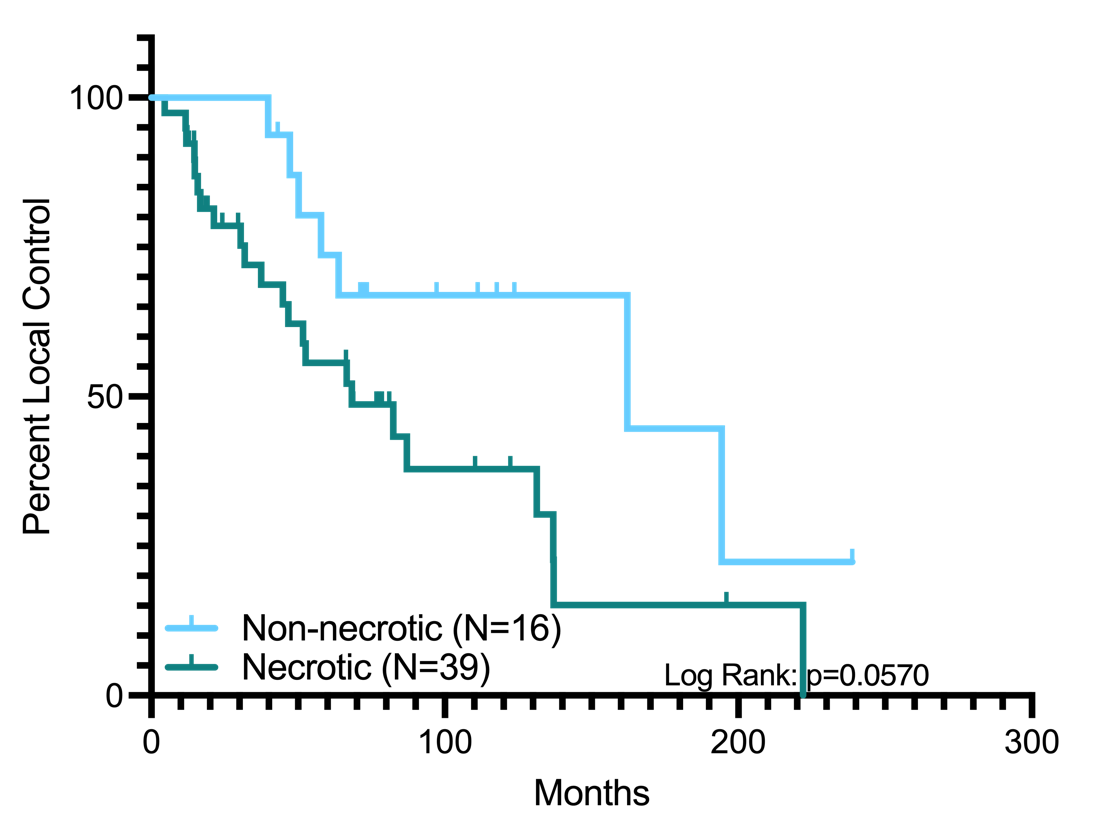

**Supplemental Figure 1. Local control based on necrosis status.**

Percent local control curve comparing post-radiation recurrence of non-necrotic G2Ms (n=16) against necrotic G2Ms (n=39). There was no significant difference between curves on log rank test (p=0.0570).

Supplemental Figure 2.

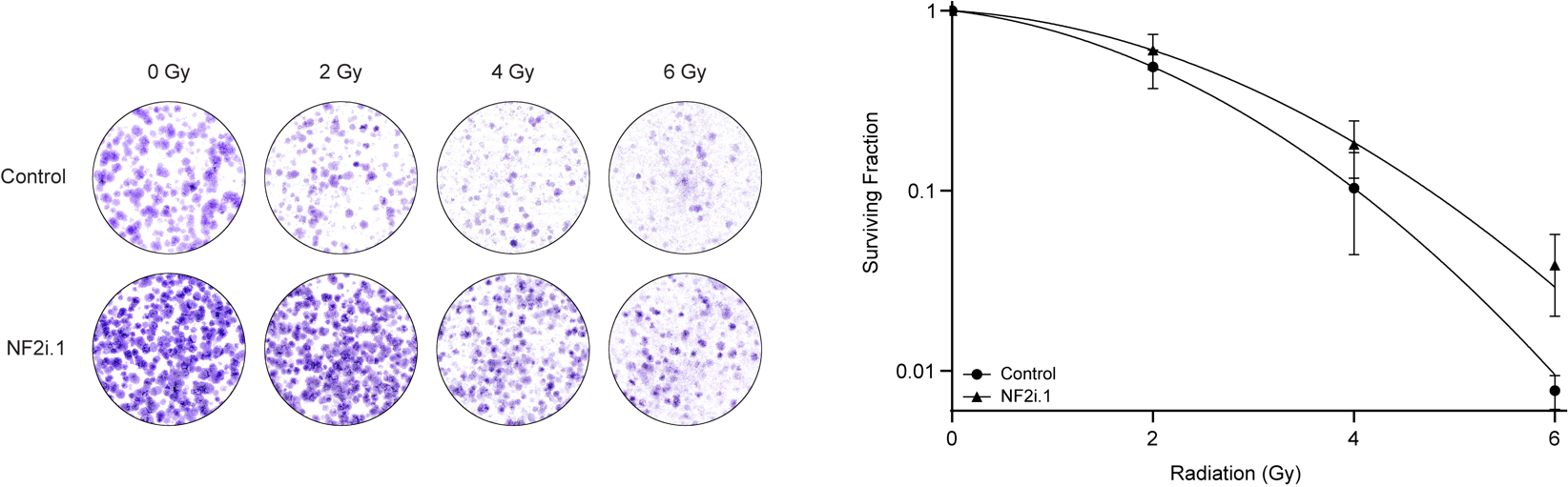
**Supplemental Figure 2. Clonogenic Assay in Hypoxia Chamber.**

Clonogenic assay with increasing doses of radiation in control, NF2i.1, and NF2i.2 IOMM Lee cells in a hypoxia chamber. There was a significant, but small difference in clonogenicity between control and NF2i.1 knockdown conditions (p=0.0010, F=10.14).

Supplemental Figure 3

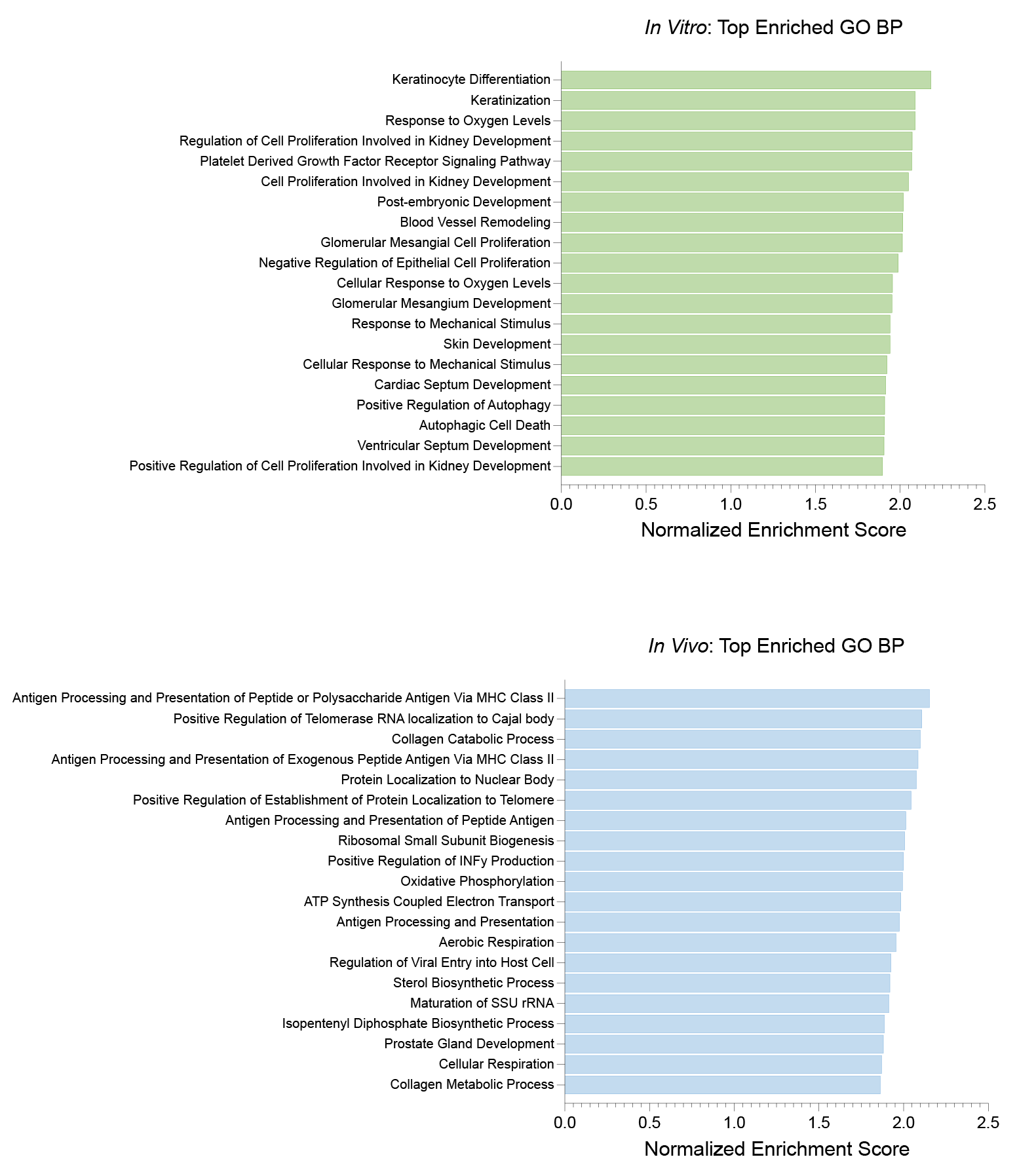

**Supplemental Figure 3. Top GO BP terms for *in vitro* and *in vivo* datasets**

Top 20 GO BP terms for both *in vitro* (*NF2* knockdown in hypoxia *vs* WT in normoxia) and *in vivo (NF2* mutation and necrosis *vs NF2* WT and non-necrotic) RNA sequencing comparisons.
